# Rapid Automated Isolation and Concentration of Bacteria from Blood Samples

**DOI:** 10.1101/2025.03.14.643371

**Authors:** Mohammad Osaid, M. Henar Marino Miguélez, Berke Baryak, Büsra Betül Özmen-Capin, Volkan Özenci, Wouter van der Wijngaart

**Affiliations:** Micro and Nanosystems, KTH Royal Institute of Technology, Stockholm, Sweden; Department of Clinical Microbiology, Karolinska University Hospital, Huddinge, Sweden

## Abstract

Sepsis is a major global health challenge associated with high mortality rates, with approximately 50 million cases yearly and 13 million deaths. For every hour a patient with septic shock remains untreated, survival decreases by 8%. Challenges persist in the rapid isolation of bacterial pathogens from blood, leading to delays in microorganism identification and antimicrobial susceptibility testing. In this study, we report a centrifuge-based device for the automated isolation and concentration of bacteria from blood samples within 40 min. The device starts with 3 ml of blood or 7.5 ml of blood culture and yields 0.7 mL of optically clear sample with a more than three-fold increased bacterial concentration, rejecting 99.97 ± 0.01% of red blood cells, 97 ± 1% of white blood cells, and 94 ± 4% of platelets. We successfully demonstrated how this process prepares blood culture samples for subsequent: sub-culturing from as low as 10 CFU/mL initial bacterial concentration; matrix-assisted laser desorption ionization time-of-flight mass spectrometry (MALDI-TOF-MS)-based identification from 5 × 10^8^ CFU/ml initial bacterial concentration; and microtrap-based detection within 1 h from 5 × 10^4^ CFU/ml initial bacterial concentration. These results demonstrate the potential of this device to automate and accelerate the diagnostic pipeline for sepsis, enabling faster detection and identification of the causative microorganism.

## Introduction

Bloodstream infections (BSIs) present a significant challenge due to delayed diagnosis and antimicrobial resistance, which can lead to sepsis. Sepsis, characterized by organ dysfunction due to a dysregulated immune response to an infection, is responsible for 20% of global deaths^1–3^. Due to the severity of this infection, patients with sepsis often require intensive care unit treatment, which incurs high costs^4^. If left untreated, it can lead to septic shock, where the mortality rate for septic patients increases by 8% per hours^5^, underscoring the critical need for rapid diagnosis in the management of sepsis. Broad-spectrum antibiotics are commonly prescribed at the onset of infection. However, this approach is suboptimal. Although it is effective in 65–80% of the cases, its failure increases fatality rates^6^ and contributes to antimicrobial resistance^7^, posing a significant threat to public health. Current state-of-the-art methods for identifying pathogens and performing antimicrobial susceptibility testing (AST) take, in general, 2-3 days^8^. This process is lengthy because it relies on multiple culture steps for identification and AST.

In clinical settings, patients’ blood samples are first subjected to a blood culture to increase the bacterial count, which typically takes between 4–72 h to produce a positive result^9–11^: approximately 50% of bottles signal positive after 16.5 h of incubation; 90% after 38.5 h, and; the mean time to positivity is approximately 21 h^12^. The time to positivity also varies based on whether the sample is from adults or neonates and the growth conditions (aerobic or anaerobic)^12^. This blood culture step is critical as only 25-38% of blood cultures test positive even in septic patients^13–15^. Once a culture is confirmed positive, it is essential to identify the microorganism and perform AST to prescribe effective antibiotics.

Current bacterial species identification methods include genotypic techniques (e.g., polymerase chain reaction), phenotypic methods (e.g., subcultures), and mass spectrometry, like MALDI-TOF, which is one of the fastest and most commonly used methods in well-equipped clinics^16^. The most common method for performing MALDI-TOF from a positive blood culture involves subculturing on an agar plate to obtain pure cultures, which adds an additional 6-12 h delay to the initiation of appropriate antibiotic treatment by another day^17^. Alternative methods, such as differential centrifugation with lysing or commercial kits like Sepsityper^18,19^ and the Accelerate Arc System^20^, are faster, but also more labor-intensive than subculturing. Traditional AST methods involve growing bacteria in broth or on agar plates in the presence of antibiotics, a time-consuming process that can take 18-24 h due to reliance on macroscopic bacterial growth identification^21^. Alternative microfluidic-based technologies can perform AST within hours by trapping and imaging single bacterial cells, but they mostly require a pure bacterial culture^22,23^. Thus, sample preparation remains a bottleneck for these technologies to function directly from blood or positive blood cultures.

Robust sample preparation that eliminates the need for culture or subculture could significantly accelerate the diagnostic process. Numerous methods and devices have been developed for sample preparation from blood or blood cultures, tailored to different downstream processing methods^16^. Common techniques for isolating and concentrating bacterial cells include sedimentation velocity-based methods^24–26^, filtration^27^, magnetic bead capture^28^, acoustophoretic techniques^29^, inertial and elastoinertial microfluidics^30,31^, surface acoustic wave (SAW)-based approaches^32^, and dielectrophoresis^33^. While centrifugation and filtration methods offer high throughput and are readily used in the clinics for sample preparation for MALDI-TOF or AST^34,35^, they require multiple steps to prepare samples for downstream processing, making them labor-intensive and susceptible to contamination. Other methods suffer from low throughput or are too complex for clinical application.

This study developed a high-throughput device for sample preparation that is non-selective, fully automated, and easy to integrate into existing diagnostic pipelines, such as MALDI-TOF, single-cell detection, and subculturing.

## Results

### Device design

We developed a centrifuge-based device for the automated sample preparation of blood samples from septic patients, designed to facilitate the downstream microbiological assays of (1) bacterial subculturing, (2) MALDI-TOF-based bacterial identification, or (3) microtrap-based bacterial detection. The device has the form factor of a standard 50 mL centrifuge tube and consists of a top and a bottom chamber connected by a siphon (Figure 1 A). The top chamber contains a grid, a cup-like structure, and two blood cell collection pockets. The bottom chamber features a rubber stop in its base. Bacterial separation and concentration are achieved in a single centrifugation protocol by concatenating velocity-based sedimentation of blood cells, automated transport of supernatant with bacteria, and sedimentation of bacteria in supernatant.

**Figure 1.**
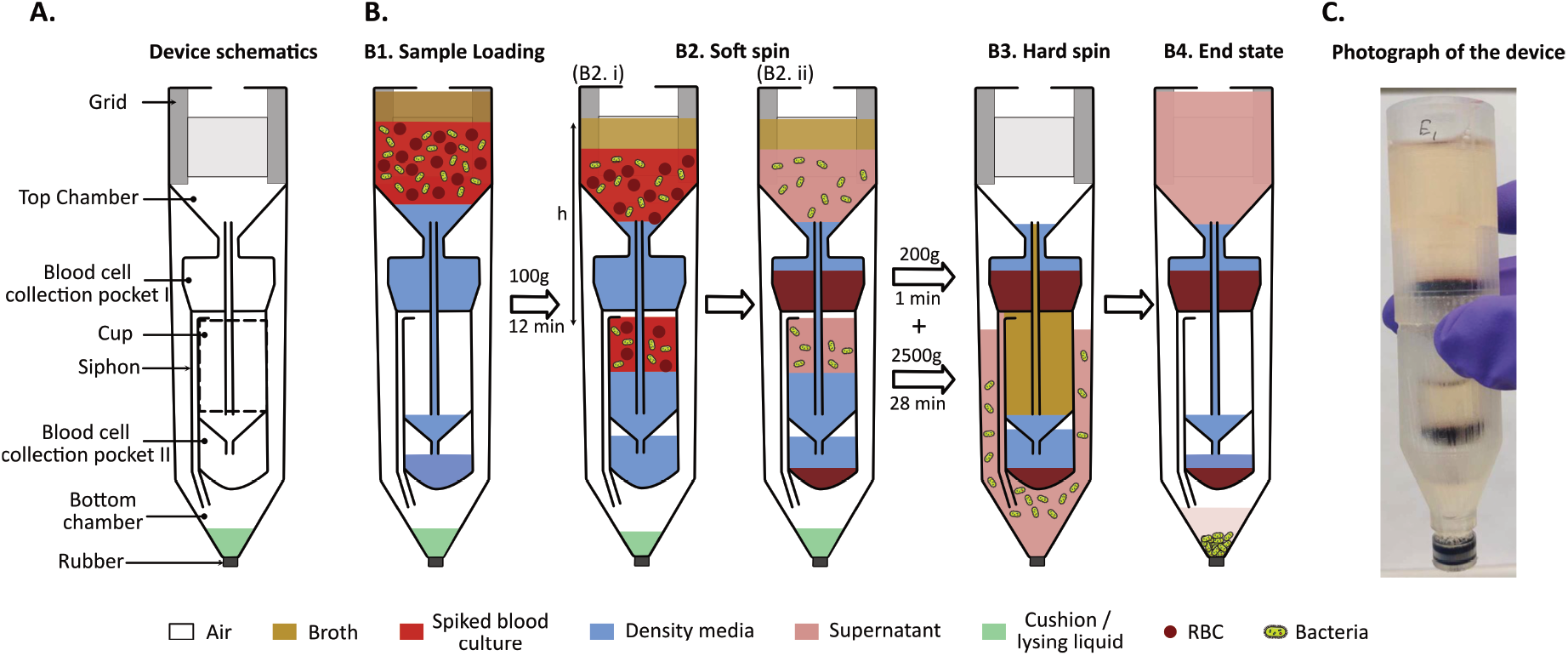
Rapid and automated isolation of bacteria from blood with a centrifuge-based device. **A.** Cross-sectional schematic of the device. The design of the device with different design features is highlighted, and the bottom chamber is pre-filled with cushion/lysing liquid. **B**. Operation of the device. **B1**. Sample loading. The top chamber was first filled with density media, and the spiked blood mixed with culture media was placed on top of it, followed by the addition of broth media. **B2**. Soft spin. The device is centrifuged at 100g to settle the blood cells in the collection pocket while preventing liquid transport to the bottom chamber. **B3**. Hard spin. The device is centrifuged at high g to move the supernatant to the bottom chamber and sediment the remaining blood cells and bacteria. **B4**. End state. The centrifuge is stopped to transfer the liquid back from the bottom to the top chamber, leaving a small volume of liquid in the bottom chamber. **C**. Photograph of a device after bacterial separation.

### Device preparation

The devices were 3D printed. The bottom chamber was pre-filled with 0.4 mL of either a lysing solution or a cushion liquid, depending on the intended downstream application. The top chamber was pre-filled with 5 mL of a density medium composed of a mixture of Lymphoprep and blood culture medium (BCM) in a ratio of 3:5. Air trapped in the cup and bottom chamber prevents the density medium from flowing down to the bottom chamber. 7.5 ml blood culture sample, consisting of 3 mL spiked blood and 4.5 mL BCM, was layered on top of the density medium. 5 mL broth medium was layered on top of the sample.

### Centrifuge operation

For a constant centrifugal acceleration, *g*^*\**^, and fluid density, *ρ*, the pressure in the trapped air in the bottom, *P*_*b*_, relates to the hydrostatic height difference in the device, *h*, as

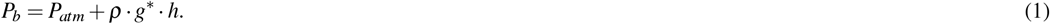

Our centrifugation protocol proceeds with a first soft spin step followed by a second hard spin step, as shown in Figure 1. The soft spin at 100 g for 12 min moves fluid into the cup and compresses the trapped air, such that the fluid fills the cup but not the bottom chamber. Blood cells exhibit terminal sedimentation velocities 20-30 times higher than bacterial cells^24^, resulting in the blood cells sedimenting into the blood collection pockets while the bacteria remain in the supernatant.

During the hard spin, the device is first accelerated to 200g for 1 min and then 2500g for 28 min. Increasing the spin speed above a critical centrifugal acceleration, *g*_*crit*_, initiates fluid transfer from the cup to the bottom chamber. *g*_*crit*_ can be designed for the range 100g < *g*_*crit*_ < 200g with an appropriate device geometry (cup volume, *V*_*c*_, and bottom chamber volume, *V*_*b*_) resulting in a hydrostatic height, *h*, for which

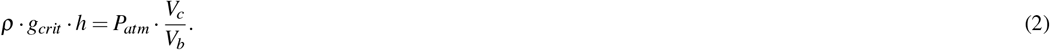

The centrifugation at 200g gently transfers supernatant to the bottom chamber without stirring the blood cell sediment in the collection pockets. The broth medium fills the cup, ensuring that all bacteria-containing liquid is transferred to the bottom chamber. During liquid transfer to the bottom chamber, the trapped air is further compressed until a new equilibrium is obtained between the hydrostatic pressure and the pressure in the compressed air volume. We verified that the liquid volume transported to the bottom chamber was consistently in the range between 8.6 and 10 mL (see SI for details). Increasing the centrifugation to 2500g facilitates the efficient sedimentation of bacterial cells in the bottom chamber. When the centrifuge stops, air decompression in the bottom chamber drives all the liquid above the siphon tube back to the top chamber, leaving the sedimented bacteria in a strongly reduced liquid volume in the bottom chamber, separated from the cup by air in the siphon. The device was then vortexed to resuspend the bacteria in the bottom chamber, after which the liquid was sampled from the bottom chamber using a syringe that pierced through the rubber stopper.

### Concatenating bacterial isolation with subculturing

Bacterial subculturing following isolation is a key step for obtaining pure colonies, enabling accurate identification and AST. To assess the potential of our centrifugal isolation method, we evaluated its performance in combination with subsequent bacterial subculturing (Figure 2). The bottom chamber was prefilled with 0.4 mL cushion fluid. The above-described centrifugation protocol resulted in approximately 0.7 mL of up-concentrated bacteria in the bottom chamber. The entire device was vortexed to resuspend the bacteria in the bottom chamber. The sample was extracted with a syringe, plated on agar plates, and left for overnight culture. The recovery rates, i.e., the number of colonies on the subculture plates relative to the number of colonies in the spiked blood sample, were 33 ± 3% (n=3), 30 ± 2% (n=3), and 48 ± 13% (n=3) for concentrations of 500, 100, and 10 CFU/mL, respectively, of *E. coli*. The recovery rates for *K. pneumoniae* and *E. faecalis* at concentration of 500 CFU/mL were 31 *±* 2% (n=3) and 34 *±* 4% (n=3), respectively. The difference in recovery is significant only between *E. coli* and *K. pneumoniae*; other differences are not significant. The rejection percentages of blood cells are 99.97 *±* 0.01% (n=4) for red blood cells, 97 *±* 1% (n=4) for white blood cells, and 94 *±* 4% (n=4) for platelets.

**Figure 2.**
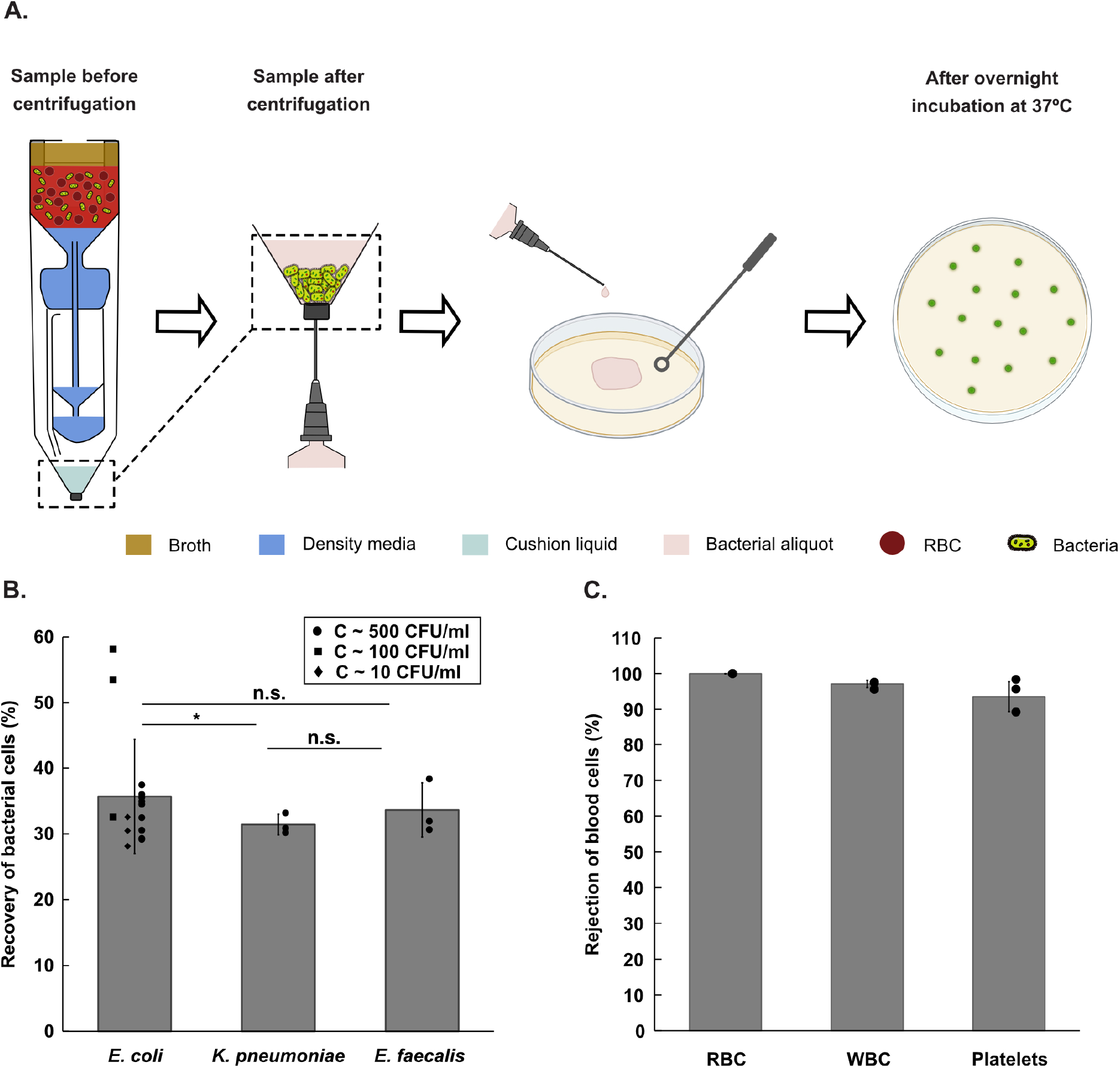
Centrifugal isolation concatenated with subculture. **A.** Centrifugal process flow for subculture directly from blood samples. **B**. Recovery of bacterial cells, being the number of colony-forming bacteria in the bottom sample after centrifugation, relative to the initial number of colony-forming bacteria in the blood sample. **C**. Rejection of blood cells, being the number of blood cells in the top and cup after centrifugation, relative to the initial number of blood cells in the blood sample. Error bars are sd; p-Value (tails 2, type 2) indicates significance: ns is not significant, * is p < 0.1.

### Concatenating bacterial isolation with MALDI-TOF-based identification

We evaluated the potential of our new isolation approach for MALDI-TOF sample preparation (Figure 3 A). The bottom chamber was prefilled with 0.4 mL of cushion fluid. A volume of 3 mL of blood was mixed with 4.5 mL of overnight bacterial culture, with each bacterial species (*E. coli, K. pneumoniae*, and *P. piersonii*) tested separately. The resulting sample for each test had an initial bacterial concentration of approximately 5 × 10^8^ CFU/mL. The 0.7 mL sample retrieved after centrifugation was mixed with 0.5 mL of deionized (DI) water and incubated at 37°C for 10 min. The mixture was centrifuged at 4000g for 5 min, and the resulting pellet was resuspended in 150 µL of solution. A 1 µL aliquot was transferred onto a MALDI target plate and air-dried at room temperature, followed by the application of the matrix solution and another air-dry, followed by analysis using a MALDI-TOF MS system. The identification score, or log score, was calculated by the MALDI-TOF MS machine based on the match between the sample’s spectrum and reference spectra. *E. coli* and *K. pneumoniae* were identified with high confidence, and the *P. piersonii* with low confidence (n = 4) as shown in Figure 3 b.

**Figure 3.**
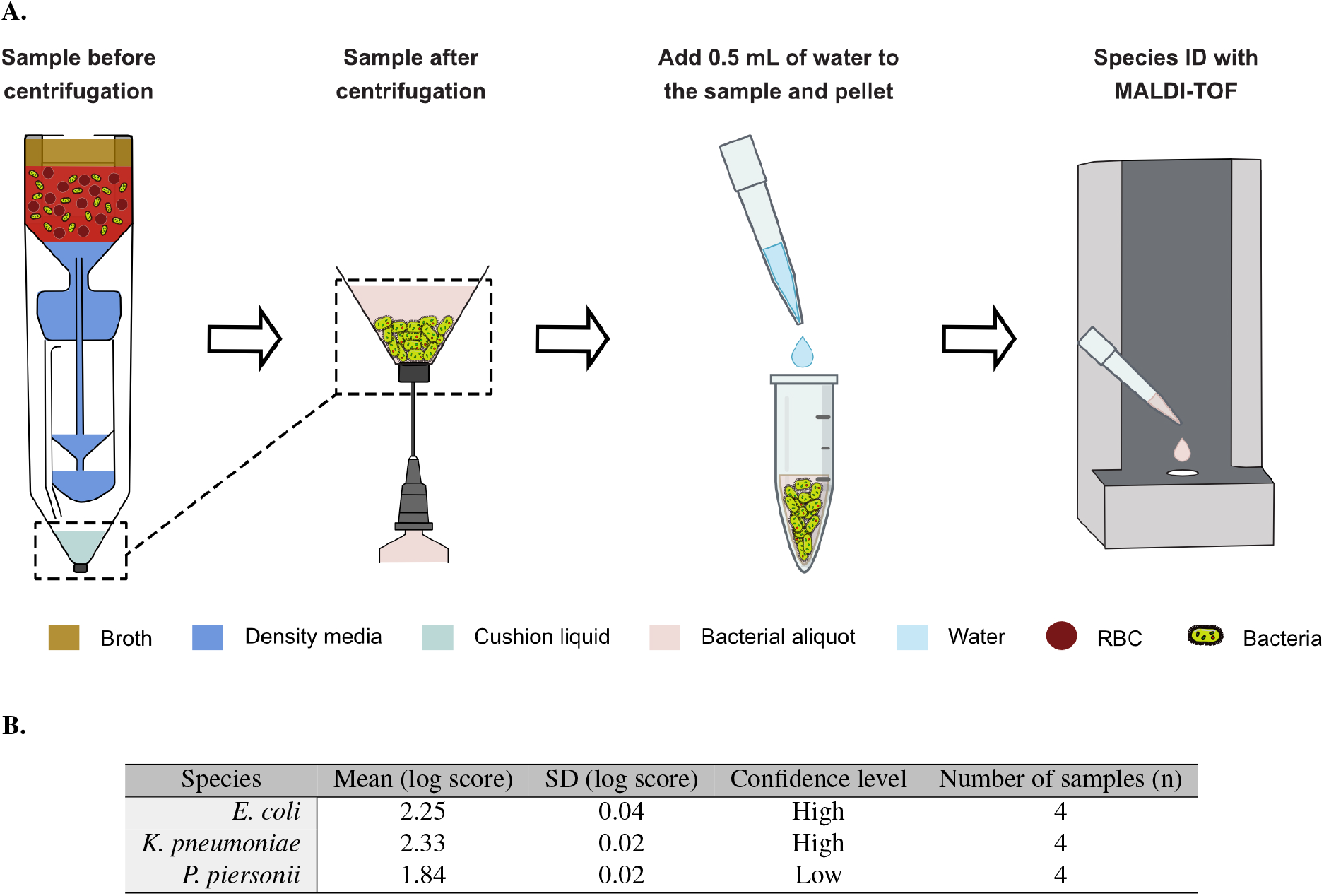
Centrifugal isolation concatenated with MALDI-TOF. **A.** Centrifugal process flow for sample preparation for MALDI-TOF. **B**. MALDI-TOF scores for three bacterial species (*E. coli, K. pneumoniae* and *P. piersonii*). Scores between 2 and 3 indicate high-confidence identification; scores between 1.7 and 1.99 indicate low-confidence identification; and scores below 1.7 represent failure to identify any species.

### Concatenating bacterial isolation with microtrap-based bacteria detection

Microtrap-based detection is a rapid and effective method for detecting bacteria directly from blood; however, it requires the efficient removal of blood cells and the concentration of bacteria^25^. To evaluate the potential of our new isolation approach for microtrap-based detection, we pre-filled the bottom chamber with 0.4 mL of lysing solution dissolved in Percoll in a 5:3 volumetric ratio (Figure 4A). Blood samples spiked with three different concentrations (5 × 10^6^, 5 × 10^5^, and 5 × 10^4^ CFU/ml) of Green Fluorescent Protein (GFP)-labeled *E. coli* were used. During hard spin centrifugation, supernatant liquid in the bottom chamber remained on top of the denser lysing solution. The centrifugation process sedimented the bacterial cells and remaining blood cells into the lysing solution, resulting in the selective lysis of the blood cells. Notably, Percoll, a density gradient medium, forms a density gradient during centrifugation, while the lysing solution remains uniformly distributed due to proper mixing at the outset. The 0.7 mL sample retrieved after centrifugation was resuspended, incubated for 10 min, and loaded into a previously developed microtrap platform^22^. For all experiments, *E. coli* cells were successfully detected using fluorescent microscopy.

**Figure 4.**
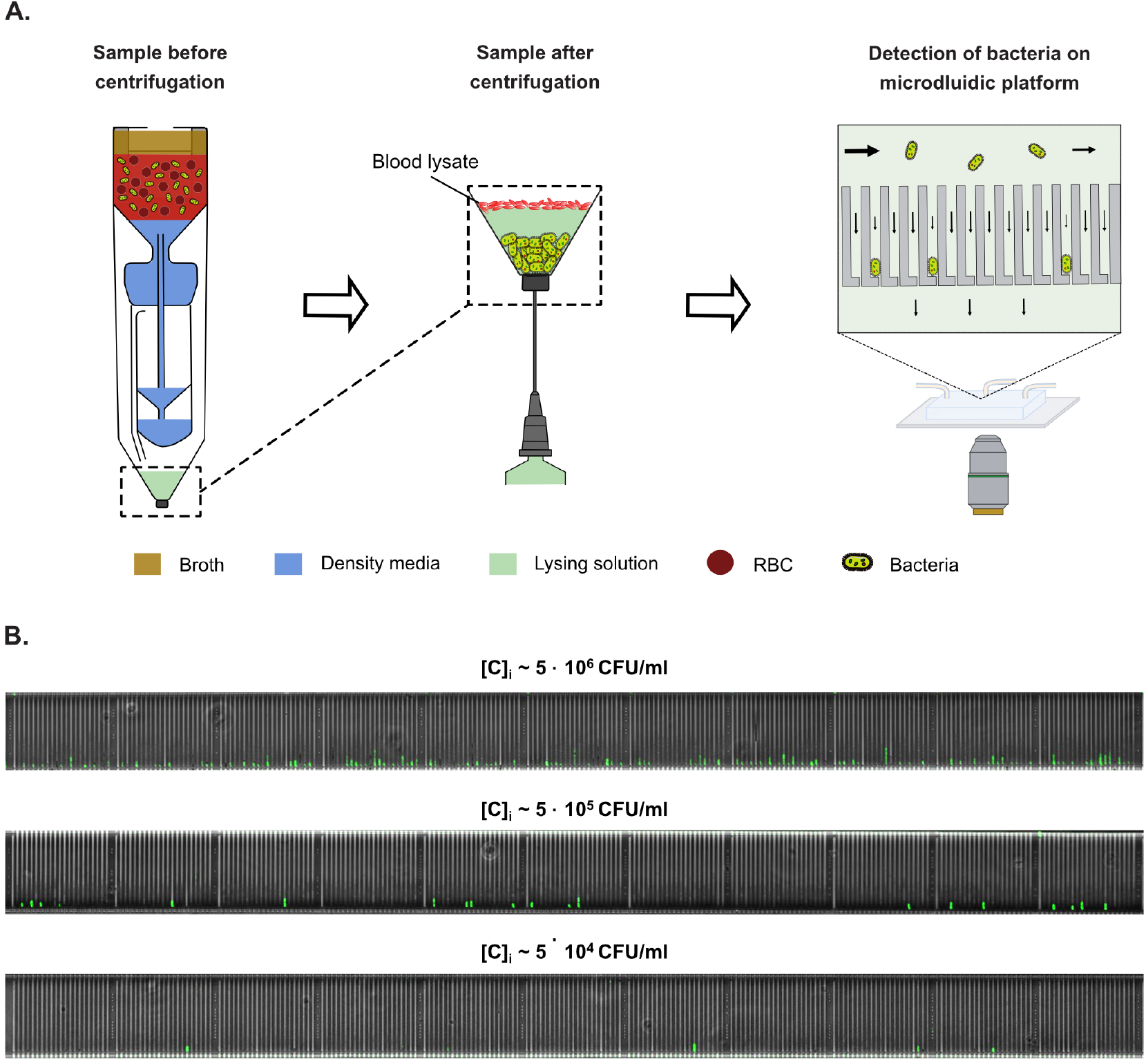
Centrifugal isolation concatenated with microtrap-based detection. **A.** Centrifugal process flow for sample preparation for microtrap-based detection. The bottom chamber of the device was prefilled with lysing solution. After centrifugation, the final liquid was sampled from the device and loaded onto a microfluidic chip. The solid arrows represent flow velocity. **B**. Fluorescence microscopy images of the microtraps with GFP-labeled *E. coli* trapped in the microtraps with three different concentrations of spiked bacteria: 5 *×* 10^6^, 5 *×* 10^5^, and 5 *×* 10^4^ CFU/ml.

## Discussion

We developed a centrifuge-based device for the automated high-throughput isolation and concentration of bacteria from blood. Combining this sample preparation with downstream assays allows for faster, less labour-intensive, and low-contamination-risk bacterial detection and identification compared to state-of-the-art workflows.

### Device design and operation

The siphon geometry in the device and the resulting air compression allow controlling the transport and metering of liquid volumes by varying the centrifugation parameters. Previously, air-compression-based valves have been used in lab-on-disk systems^36^; however, these systems were limited in throughput, required non-standard laboratory equipment, and needed active valves. Our device addresses these limitations by handling 7.5 mL of blood culture and operating on standard centrifuges without additional valves. During device design, we needed to solve specific challenges related to the relatively large volume and device size, specifically vacuum boiling, blood cell resuspension during fluid transport, and large dead volumes. These challenges were addressed through the careful selection of design parameters for the cup, siphon, blood cell collection pockets, and grid, as detailed below.

#### Design and position of the cup and siphon

The cup constitutes an internal fluid volume that allows partial compression of the trapped air without liquid transfer to the bottom chamber during the soft spin. The siphon entry is placed at the top of the cup, such that the entire air volume in the cup, *V*_*c*_, can be used for compression. The cup is positioned such that its entire lumen remains below the top liquid surface at any time during operation, thus avoiding negative hydrostatic pressure leading to liquid boiling (details in Supplementary Information). Increasing the cup volume allows increasing the relative centrifugal force (RCF) during the soft spin without transferring liquid to the bottom chamber, as described in the equation 2. A large cup volume, however, forms a dead volume for the liquid transferred from the top chamber to the bottom chamber, risking the trapping of bacteria in the cup during their intended transfer to the bottom chamber. To mitigate the latter, a layer of broth is added above the spiked blood mixture such that during hard spin, the broth fills the entire lumen of the cup, thereby driving the bacteria-rich supernatant into the bottom chamber.

#### Blood cell collection pocket design

The connections to the blood cell collection pockets were kept narrow to reduce the swirling of blood cells during centrifugal acceleration or deceleration, hence avoiding unwanted resuspension and blood cell transfer to the bottom chamber.

#### Grid design

To prevent the broth from mixing with the supernatant during low-speed to high-speed centrifugation acceleration, a rectangular grid-like structure was introduced at the top of the top chamber^27^. Additionally, the cross-sectional area of the lower part of the top chamber was reduced. These modifications minimize the effects of Coriolis and Euler forces, which could otherwise mix the sample and reduce the efficiency of velocity-based sedimentation.

#### Preloaded liquids for separation

To isolate bacteria from blood, the spiked blood mixture was layered over density media, facilitating separation based on velocity differentiation. The falling distance through the density media enhances the separation of bacteria from blood cells. Further, a cushion liquid in the bottom chamber minimizes bacterial loss due to excessive centrifugation.

#### Liquid transfer analysis

In all the above experiments, the top chamber was filled with 17.5 mL of liquid. During the hard spin, between 8.6 and 10 mL of liquid was transferred to the bottom chamber, while 8–9 mL remained in the top chamber and cup (Supplementary Figure 2). Of the liquid remaining in the top chamber, approximately 4 mL was retained in the blood cell collection pockets, approximately 4 mL in the cup, and the rest constituted the dead volume above the blood cell collection pockets. The liquid retained in the blood cell collection pockets contained mostly blood cells, whereas the cup-like structure was filled with broth media, which should contain few bacterial cells as it remains at the top of the blood mixture.

### Comparison with state-of-the-art bacterial isolation

Current methods for isolating and concentrating bacteria from blood have significant limitations, including low up-concentration factors^24,25,27,29–32^ (the ratio of bacterial concentration in the final aliquot to the initial sample) or limited throughput^27–33^ (Supplementary Table 1). These challenges hinder their clinical use, as diagnosing sepsis or BSIs requires addressing extremely low bacterial concentrations in clinical samples. Our approach addresses these challenges with an up-concentration factor above three and a throughput of 75 µL/min, outperforming most existing methods. In contrast to many methods that work well only at higher bacterial concentrations^29,30,32,33^, our approach works reliably even at 10 CFU/mL. While most methods focus only on bacterial isolation, ours performs full sample preparation, including bacterial concentration and selective cell lysis. Additionally, our device operation is scalable, with up to 16 units operating simultaneously in centrifuges equipped with 16 holders, such as the Thermo Scientific™ Sorvall™ ST 16 Centrifuge, thus supporting high-throughput processing. These benefits make our approach a practical and efficient solution for clinical diagnostics.

**Table 1.** Key performance parameters of smart centrifugation and other separation methods.

|  | Up-concentration factor | Minimum concentration detected (CFU/ml) | Blood processed (ml) | Time (min) | Throughput (μl/min) | RBC removal efficiency |
| --- | --- | --- | --- | --- | --- | --- |
| This device | 3.20 | 14 | 3 | 40 | 75 | 99.97% |
| Smart centrifugation | 0.58 | 9 | 2.25 | 5 | 450 | 99.98% |
| Filter based centrifugation<sup>27</sup> | 0.01 - 0.02 | 10 | 1 | 60 | 16.7 | 99.4% |
| Compact Disk<sup>24</sup> | 0.67 | 10 | 7 | 1 | 7000 | 94% |
| Dextran sedimentation<sup>26</sup> | 1.25-1.50 | 10-100 | 10 | 30 | 333 | >90% |
| Elastoinertial Separation<sup>31</sup> | 0.43 | 40 | 1 | 40 | 25 | 100% |
| Inertial lift forces<sup>30</sup> | 0.02 | >10<sup>5</sup> | 0.15 | 4 | 37.5 | 90% |
| Dielectrophoresis<sup>33</sup> | 87.20 | 10<sup>4</sup> | 0.05 | 75 | 0.67 | 100% |
| SAW<sup>32</sup> | 0.035 | 4.4.10<sup>7</sup> | 0.012 | 600 | 0.02 | 99.6% |
| Acoustophoresis<sup>29</sup> | 0.25 | 5.10<sup>8</sup> | 1 | 50 | 20 | 99.99% |
| Magnetic Bead Separation<sup>28</sup> | 4.00 | 1 | 5 | 60 | 83.3 | >99.99% |

### Preparation for bacterial subculture

Subculturing is a method for obtaining pure solid bacterial colonies, which are required for MALDI-TOF and genotypic-based detection. Our method prepares blood samples for subculturing in less than 1 h, even at bacterial concentrations as low as 10 CFU/mL, reducing the procedure time by hours to days compared to the current praxis requiring *a priori* positive blood cultures (Figure 5).

**Figure 5.**
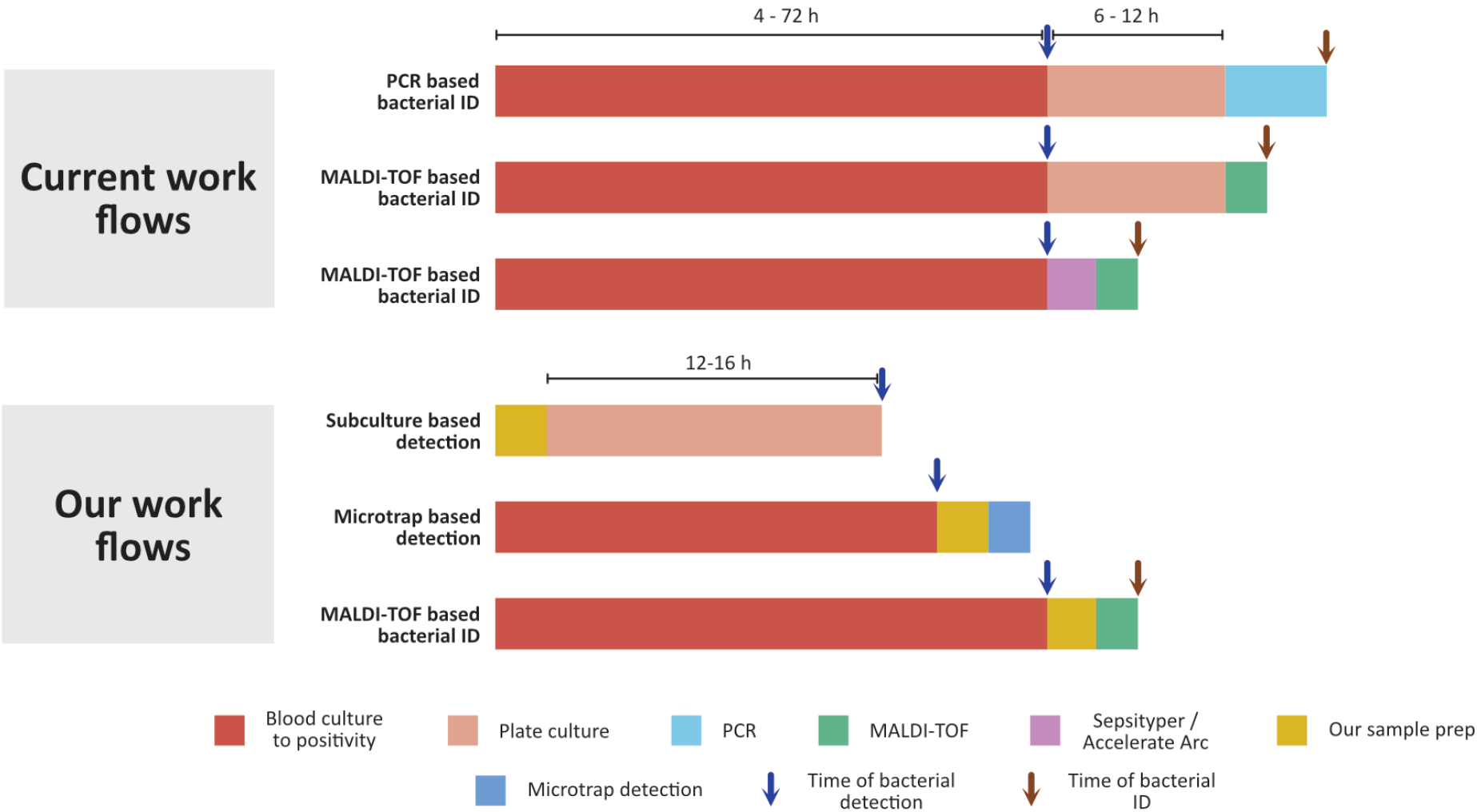
Comparison of the State-of-the-Art Workflow vs. Our Sample Preparation Device. Current workflows for bacterial identification typically involve blood culture followed by plate culture and PCR-based identification or the MALDI-TOF workflow, which also requires blood culture followed by plate culture or the use of commercially available kits such as Sepsityper or Accelerate Arc Systems for identification. In our subculture-based detection method, bacterial detection is performed without blood culture by utilizing our sample preparation device, followed by overnight incubation. For Microtrap-based detection, a high bacterial concentration is not required, allowing detection before the culture turns positive. In MALDI-TOF-based detection, we start with a high bacterial concentration, similar to that of a positive blood culture, and identify bacteria directly after our sample preparation, eliminating the need for plate culture.

### Preparation for MALDI-TOF-based identification

We successfully demonstrated the suitability of our device for sample preparation for MALDI-TOF analysis. In most clinical settings, standard MALDI-TOF sample preparation requires subculturing of positive blood cultures, a process that takes 6–12 h and can delay appropriate antibiotic therapy by up to a day.

In contrast, our device produces, in less than 1 h, purified bacterial aliquots directly suited for MALDI-TOF identification, enabling accurate determination of the causative bacterial species with high score values. Compared to other state-of-the-art sample preparation kits, such as the Sepsityper^18,19^ or Accelerate Arc System^19,20^, our device requires less hands-on time.

### Preparation for microtrap-based detection

We successfully demonstrated the suitability of our device for preparing samples for bacterial detection on microtrap platforms. Microfluidics-based bacterial culture is a promising technology with the potential to significantly advance diagnostic methods, particularly in its ability to perform rapid phenotypic testing by analyzing the growth of single bacterial cells, unlike traditional clinical practices that rely on macroscopic growth^22^. Additionally, bacteria can be identified on such platforms using genotypic methods such as FISH^37^. However, microfluidic devices typically require pure bacterial cultures, as interfering cells readily clog the microtraps—especially those with dimensions of just a few microns—thereby compromising the assay. Current methods and devices for sample preparation from whole blood for microfluidic-based detection are multiple-step, labour-intensive and require trained personnel.^25,26^

In contrast, our device simplifies bacterial isolation by automating multiple centrifugation steps and blood cell lysis in a single centrifugation protocol. Moreover, initial bacterial concentrations as low as 5 × 10^4^ CFU/ml yielded positive detection, which is significantly lower than the levels typically found in positive blood cultures.

## Conclusion

In conclusion, we introduce a centrifugation device capable of performing sample preparation for multiple applications in a single centrifugal operation. This innovation speeds up sample preparation and reduces its labour intensity, potentially facilitating the integration of advanced diagnostic technologies into clinical settings. By streamlining multiple preparation steps into a single automated process, our device enhances efficiency and reproducibility while minimizing the risk of cross-contamination. Its adaptability for clinical assays further underscores its potential for widespread adoption in both research and point-of-care diagnostic settings.

## Methods

### Device fabrication

The device was designed using SolidWorks CAD software and fabricated in three distinct parts: top, bottom, and cap, as detailed in the Supplementary Figure 3. Fabrication was performed using a Form 3+ 3D printer (Formlabs, USA) with Clear V4 resin. Post-fabrication, the components were cleaned using a Form Wash (Formlabs, USA) with isopropanol (IPA) to remove residual resin from the surfaces. The internal structures of the device were manually cleaned using a needle and IPA to ensure thorough removal of excess resin. Subsequently, the parts were cured in a Form Cure (Formlabs, USA) using ultraviolet (UV) light for 30 min at 60 °C. To address challenges during 3D printing, a hole was incorporated into the blood cell collection pocket II of the top part to prevent cupping. This hole was sealed post-fabrication using Clear V4 resin to prevent leakage during operation.

### Device assembly

The top and bottom 3D-printed components were adhered using Clear V4 resin, which was solidified through curing with UV light in the Form Cure for 10 min. A polyisoprene rubber plug was inserted into the cap and glued using ClearSeal Glass Clear adhesive (Casco, Switzerland), then allowed to dry overnight. Subsequently, the cap was attached to the bottom component using Clear V4 resin. Prior to this step, the bottom chamber was pre-filled with a cushion or lysing liquid according to the use case.

### Medium prepration

The density medium used in this study was a mixture of Lymphoprep (STEMCELL Technologies, Canada), a medium with a density of 1.077 g/mL, and blood culture medium (BCM) (BD BACTEC Plus Aerobic medium, BD, USA) in a 3:5 ratio. The broth was prepared by dissolving Luria low salt powder (L3397, Sigma-Aldrich, USA) in deionized water (DIW) at a concentration of 25 g/L, followed by autoclaving to ensure sterility. The cushion liquid employed was a combination of Percoll (Sigma-Aldrich, USA), a density medium with a density range of 1.125–1.135 g/mL, and BCM mixed in a ratio of 3:5 . The lysing solution used in the study was prepared by mixing Percoll with a lysing liquid in a 3:5 ratio. The lysing liquid itself consisted of 2% (w/v) sodium cholate hydrate (Sigma-Aldrich, USA) and 1% (w/v) saponin (Sigma-Aldrich, USA), both dissolved in BCM.

### Bacterial strains

Four different bacterial strains were used in this study. The *E. coli* strain carried a plasmid expressing mVenusNB fluorescence proteins. The other bacterial strains, *K. pneumoniae, E. faecalis*, and *Pantoea piersonii*, were clinical strains randomly collected from a clinical microbiology laboratory in Sweden. For long-term storage, the bacteria were maintained at -80 ^°^C in standard glycerol solution. Prior to use, the strains were incubated overnight at 37 ^°^C in BD BACTEC Plus Aerobic medium (BD, USA), referred to as Blood Culture Medium (BCM). After incubation, the bacterial cultures were diluted in BCM to approximately 10^4^, 10^3^, and 10^2^ CFU/mL, which were subsequently used to spike blood samples at varying concentrations for the experiments. Further, the concentration of the spiking solution was determined by plate counting of bacterial solution of 10^3^ CFU/ml.

### Spiked blood prepration

In the experiments, healthy donor blood was obtained from the blood bank (Blodcentralen, Stockholm, Sweden). The blood samples were used within two days of collection and stored at 4 ^°^C prior to use. The blood was diluted with BCM in a 2:3 ratio and spiked with bacteria. During spiking, the volume of the liquid added was always less than 4% of the total volume of the blood culture.

### Bacteria counting

Bacterial counting was conducted by plating the bacterial solution on agar plates, followed by overnight incubation at 37 °C. The agar plates were prepared by dissolving LB broth with agar (Miller) (Sigma-Aldrich, USA) in deionized water at a concentration of 40 g/L. The mixture was autoclaved, poured into Petri dishes, and cooled. For all subculture experiments, 100 µL of the final aliquot was plated in triplicates. However, the entire final aliquot was plated in the 10 CFU/mL experiments to ensure accurate quantification.

### Blood cell quantifiation

Blood cell counts in whole blood and final aliquots were measured using a hematology analyzer (Swelab Alfa Plus, Boule Diagnostics, Sweden).

### Process for MALDI-TOF

A 1 µL aliquot of the final sample was applied to the MALDI-TOF plate and air-dried at room temperature. Subsequently, the matrix solution was applied, and the sample was analyzed using the MALDI-TOF MS system after drying the matrix solution. The MALDI-TOF machine used in the study was MALDI Biotyper® Sirius One System (Bruker Daltonics, Bremen, Germany).

### Microfluidic platform fabrication

The design and the fabrication of the microfluidic chip were previously reported^22,38^. The silicon mold used for the chip was manufactured by ConScience AB, Sweden, and was silanized for 30 min before replication with polydimethylsiloxane (PDMS) (Sylgard 184, DOW, USA). A mixture of PDMS and curing agent in a 10:1 w/w ratio was poured onto the mold and cured at 80 °C overnight. Openings for the PDMS ports (2.0, 2.1, 2.2, 5.1, and 5.2; see chip design in^22,38^) were created using a 0.5 mm puncher. The fabricated PDMS stamps were cleaned with isopropanol (IPA) before being bonded to a glass coverslip (No. 1.5, Menzel-Gläser, Germany) using plasma treatment. The bonded assembly was subsequently heat-cured for an h at 80 ^°^C.

### Flow control

The microfluidic device was set up on a microscope and connected to reservoirs using tubing (TYGON, Saint-Gobain, North America). The flow controller used for pressurizing the reservoirs was the FlowEZ (Fluigent, France). For priming, the reservoirs connected to ports 2.1, 2.2, 5.1, and 5.2 were initially pressurized at 500 mbar with water containing 0.085 g/L of Pluronic F108 (Sigma-Aldrich, USA), after which the pressure was reduced to 0 mbar. Subsequently, the reservoir connected to inlet port 2.0 was pressurized at 500 mbar for priming and then increased to 1000 mbar for sample loading.

### Optical setup

Images of the microtraps were captured at 100x magnification using a Nikon Eclipse Ti-U inverted microscope. Fluorescence images were also taken at 100x magnification using a CFI Plan Fluor DLL 100x (1.30 NA, oil) objective. The fluorescence images were acquired with a filter cube comprising a FF497-Di01 dichroic mirror (Semrock, USA), a FF01-469/35 excitation filter (Semrock, USA), and a FF01-525/39 emission filter (Semrock, USA).

## Acknowledgements

This work was funded through the Swedish Foundation for Strategic Research (SSF) via the Agenda 2030 Research Centre “Ultra-rapid antibiotic susceptibility determination” ARC19-0016.

## Supplementary

### Setup to film the liquid motion in the device inside the moving centrifuge

An imaging setup was developed to monitor liquid motion within the device during centrifugation. A customized centrifuge holder was designed, 3D printed, and engineered to accommodate a camera, a light source, and the device, all of which fit inside a standard centrifuge bucket (Supplementary Figure 1). The holder incorporates an integrated wireless camera (Global Tsolar Lights Electrical, China), a light source (Ledlenser, Germany), and the fluidic device. During centrifugation, the wireless camera transmits real-time images of the process to an external computer, enabling continuous monitoring.

**Supplementary Figure 1:**
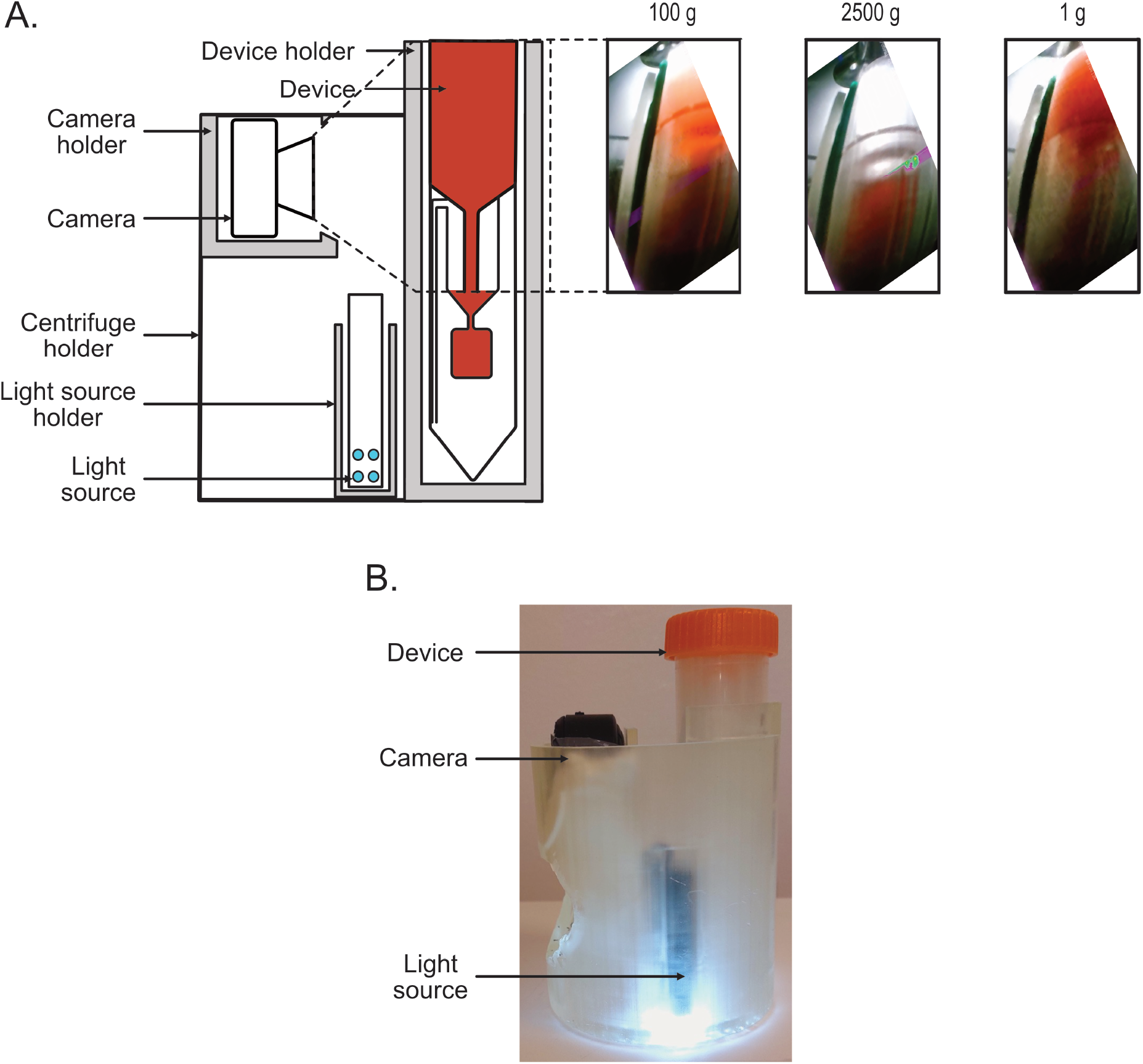
Setup for filming inside a centrifuge tube: **A**. Schematics of the holder used for filming the device inside the centrifuge, which includes compartments for the camera, the device, and the light source. On the right are the Images of liquid movement within the device during centrifugation taken from the camera. The figure includes snapshots of the device inside the centrifuge at 100g (soft spin), 2500g (hard spin), and 1g (centrifuge stopped). **B**. Photograph of the holder with the camera, the device, and the light source in place.

The device was filled with blood mixed culture media and placed in the imaging setup. During centrifugation at 100g (soft spin), it was observed that the liquid level rose inside the cup-like structure, and the liquid level in the top chamber decreased, but no liquid was transferred to the bottom chamber (Supplementary Figure 1 A). However, during hard spin centrifugation at 2500g, the liquid was transferred to the bottom chamber, leaving only a residual volume in the cup-like structure. When the centrifuge was stopped, the liquid returned to the top chamber, leaving only a small amount of liquid in the bottom chamber.The video can be found in Supplementary files. The image of the imaging system is shown in (Supplementary Figure 1 B).

### Calculation of Volume transferred to the bottom

The transfer of liquid to the bottom chamber was visualized using a camera placed inside the centrifuge tube during operation, as shown in Supplementary Figure 1. To precisely quantify the volume transferred, multiple small grooves or traps were introduced along the inner wall of the chamber. The volume below these traps, extending to the bottom of the tube, was measured manually. In other words, the bottom chamber of the tube was calibrated using these traps, as highlighted in Supplementary Figure 2.

During a hard spin, the liquid transferred to the bottom chamber filled the traps that became submerged. After filling the top chamber with 17.5 mL of liquid and centrifuging the device at 2500g, the first three traps from the bottom were filled, while the fourth trap at the top remained empty. The volumes corresponding to the regions below the third and fourth traps were 8.6 mL and 10 mL, respectively. Therefore, the volume of liquid transferred was determined to be between 8.6 mL and 10 mL.

**Supplementary Figure 2:**
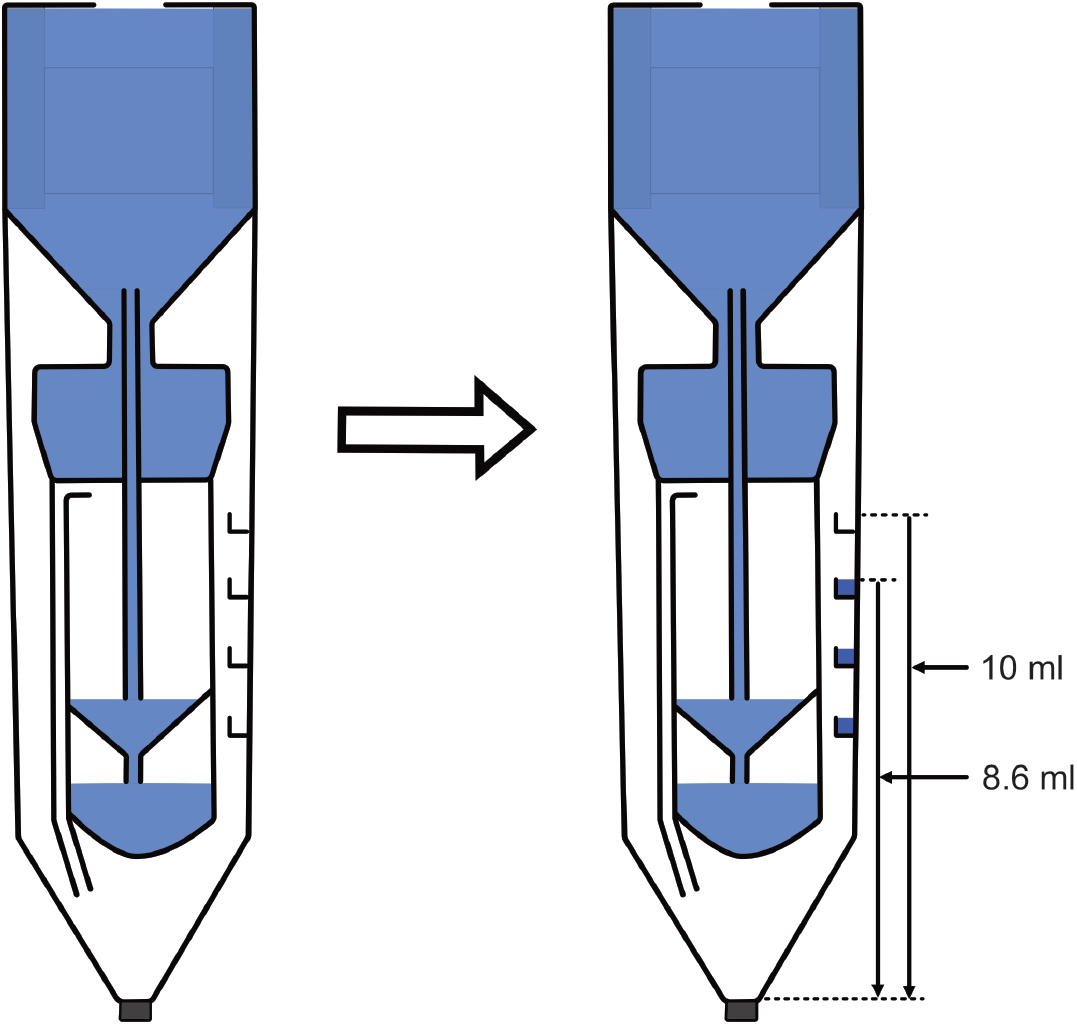
Device design for quantifying the volume of liquid transferred to the bottom chamber. Schematic of the device with microtraps in the bottom chamber. During hard spin, the liquid gets transferred to the bottom chamber and fills the microtraps. When centrifugation is stopped, the liquid returns to the top chamber, leaving the traps filled.

### Device assembly

The device consists of three distinct 3D-printed components that are assembled together. The two larger parts, namely the Top and Bottom parts, as shown in Supplementary Figure 3 A, were bonded using clear v4 resin and subsequently cured in ultraviolet light in a Form Cure system (Formlabs, USA) for 10 min. The bottom cap incorporates a polyisoprene rubber plug sourced from 3 mL Megro™ SOFT-JECT™ disposable syringes (Henke-Sass, Wolf GmbH). This plug was attached to the 3D-printed cap using ClearSeal Glass Clear adhesive (Casco, Switzerland) and allowed to dry overnight. The cap was then affixed to the device using clear v4 resin and cured in ultraviolet light for an additional 10 min. Supplementary Figure 3 B illustrates the CAD model of the assembly of the device.

**Supplementary Figure 3:**
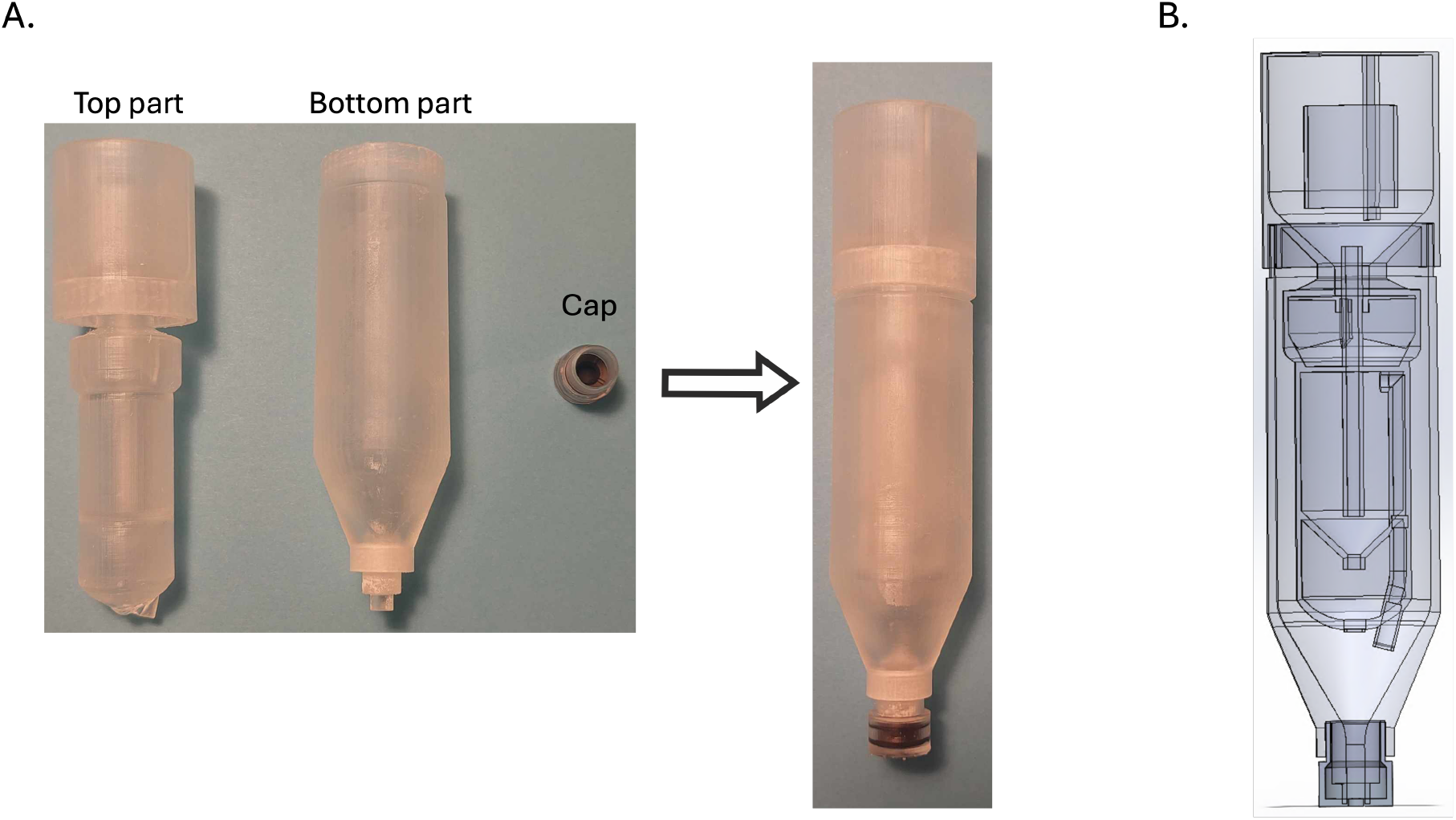
Assembly of the 3D printed device. **A**. Image showing the three separate 3D-printed components of the device, including the Top, Bottom, and cap, which are bonded using clear v4 resin and cured under ultraviolet light to form the final structure. **B**. CAD model illustrating the cross-section of the complete assembly of the device.

### Vaccum creation in the centrifugal device

The device described in the article can be modified to create a vacuum by changing the position of the cup, i.e., by moving it upward in the top chamber, as shown in Supplementary Figure 4. The top chamber of the device is filled with liquid, and during a high-speed spin (centrifugation at 2500g), the liquid is transferred to the bottom chamber via the siphon, which consists of the cup and tube. Throughout the transfer process, the liquid surface in the top chamber remains at atmospheric pressure. When the liquid level in the top chamber drops below the top edge of the cup, the pressure inside the cup is given by Equation 3:

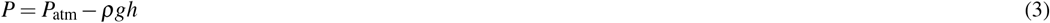

The liquid begins to boil when the pressure inside the cup, as described by the equation, approaches zero. This boiling prevents further transfer of the liquid, resulting in the state shown in Supplementary Figure 4. The video can be found in Supplementary files.

**Supplementary Figure 4:**
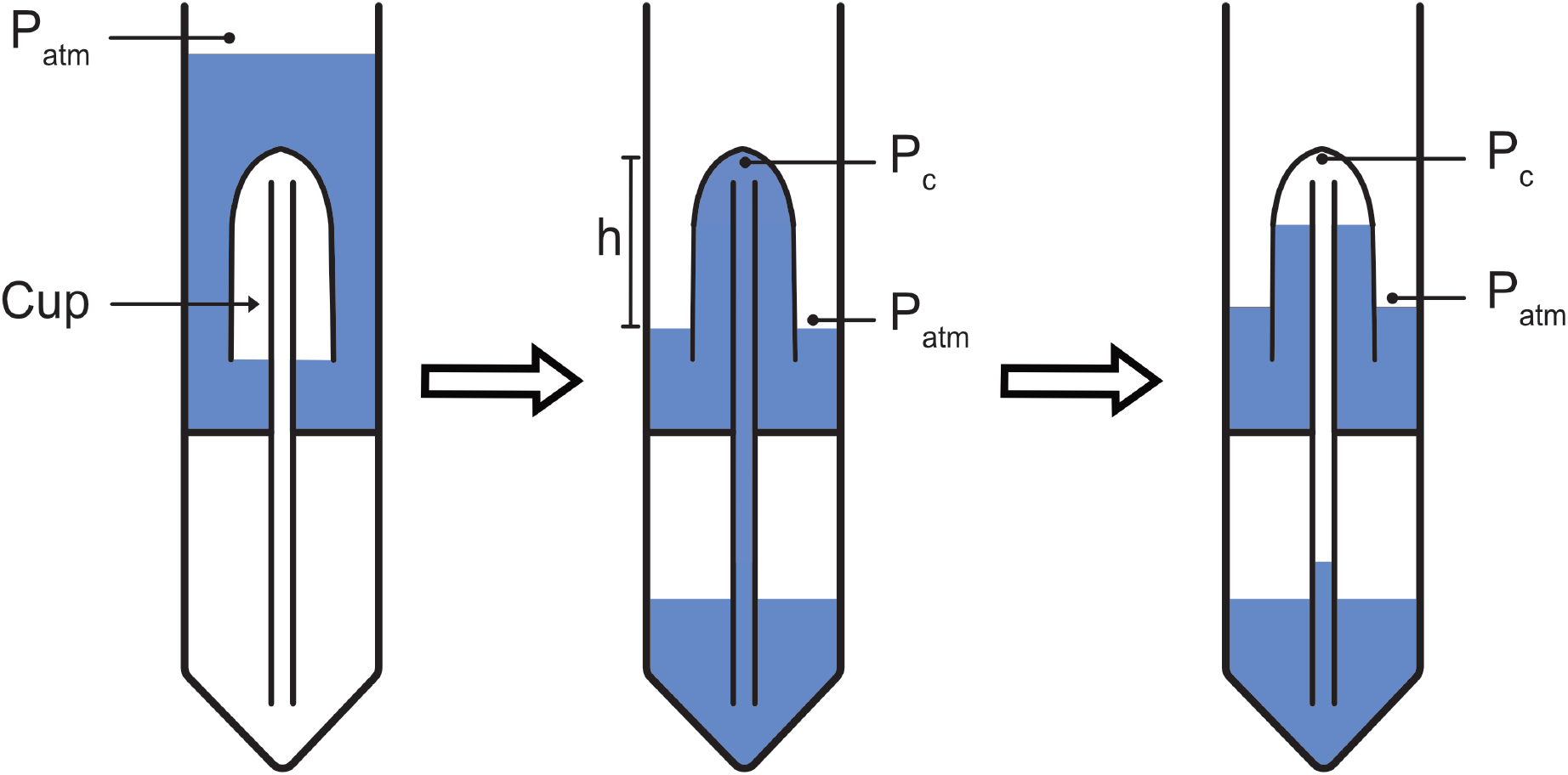
Vacuum generation in the device by adjusting the cup’s position. During high-speed centrifugation (2500g), liquid transfer occurs via the siphon. When the liquid level in the top chamber drops below the top edge of the cup, the pressure inside the cup decreases, causing the liquid to boil and preventing further transfer.

### Comparison with other separation methods

The key parameters for sample preparation methods from the blood for downstream detection include up-concentration factor, the minimum concentration of bacteria, blood processing capacity, throughput, and cell removal efficiency. These parameters are critical for the detection of bacterial presence, identification, and AST in the clinical management of sepsis and bloodstream infections (BSIs).

Up-Concentration Factor: The up-concentration factor is defined as the ratio of the bacterial concentration in the final aliquot to the initial concentration in the blood or blood culture. A higher up-concentration factor is advantageous for downstream detection, as the primary challenge in diagnosing sepsis or BSIs lies in the extremely low bacterial concentrations typically present in clinical samples. The developed device exhibits a significantly high up-concentration factor, making it well-suited for these applications.

Minimum Concentration of Bacteria: Given the low bacterial concentration in blood or blood culture during sepsis or BSI, it is crucial for any method or device to perform efficiently at such low levels. The developed device has been demonstrated to function effectively at bacterial concentrations as low as 10 CFU/mL, which is clinically relevant. This highlights its potential utility in addressing one of the most challenging aspects of sepsis diagnosis.

Throughput and Blood Volume Processing: Due to the low bacterial concentrations in clinical scenarios, processing large volumes of blood with high throughput is essential. The developed device demonstrates a superior throughput compared to many existing methods, as illustrated in the accompanying comparative data table. This capability enhances its applicability for routine clinical use.

Cell Removal Efficiency: In addition to bacterial concentration, the number of residual blood cells in the final aliquot significantly impacts the accuracy of bacterial detection. Residual blood cells can interfere with downstream detection processes, making blood cell removal efficiency a critical parameter. The developed device achieves a red blood cell (RBC) removal efficiency of 99.99%, which is exceptionally high and comparable to, if not better than, other existing methods.

In summary, the device demonstrates excellent performance across all critical parameters for sample preparation from blood or blood mixed with culture media, underscoring its potential for enhancing the detection of sepsis and bloodstream infections in clinical settings.

## References

1. Rudd, K. E. et al. Global, regional, and national sepsis incidence and mortality, 1990–2017: analysis for the global burden of disease study. The Lancet 395, 200–211 (2020).

2. Hotchkiss, R. S. et al. Sepsis and septic shock. Nat. reviews Dis. primers 2, 1–21 (2016).

3. Thompson, K., Venkatesh, B. & Finfer, S. Sepsis and septic shock: current approaches to management. Intern. medicine journal 49, 160–170 (2019).

4. Arefian, H. et al. Hospital-related cost of sepsis: a systematic review. J. Infect. 74, 107–117 (2017).

5. Kumar, A. et al. Duration of hypotension before initiation of effective antimicrobial therapy is the critical determinant of survival in human septic shock. Critical care medicine 34, 1589–1596 (2006).

6. Rhee, C. et al. Prevalence, underlying causes, and preventability of sepsis-associated mortality in us acute care hospitals. JAMA network open 2, e187571–e187571 (2019).

7. Llor, C. & Bjerrum, L. Antimicrobial resistance: risk associated with antibiotic overuse and initiatives to reduce the problem. Ther. advances drug safety 5, 229–241 (2014).

8. Morgenthaler, N. G. & Kostrzewa, M. Rapid identification of pathogens in positive blood culture of patients with sepsis: review and meta-analysis of the performance of the sepsityper kit. Int. journal microbiology 2015, 827416 (2015).

9. Jacobs, M. R. et al. Multicenter clinical evaluation of bact/alert virtuo blood culture system. J. Clin. Microbiol. 55, 2413–2421 (2017).

10. Lambregts, M. M., Bernards, A. T., van der Beek, M. T., Visser, L. G. & de Boer, M. G. Time to positivity of blood cultures supports early re-evaluation of empiric broad-spectrum antimicrobial therapy. PLoS One 14, e0208819 (2019).

11. Kumar, Y., Qunibi, M., Neal, T. & Yoxall, C. Time to positivity of neonatal blood cultures. Arch. Dis. Childhood-Fetal Neonatal Ed. 85, F182–F186 (2001).

12. Ransom, E. M., Alipour, Z., Wallace, M. A. & Burnham, C.-A. D. Evaluation of optimal blood culture incubation time to maximize clinically relevant results from a contemporary blood culture instrument and media system. J. clinical microbiology 59, 10–1128 (2021).

13. Yu, D. et al. Correlation of clinical sepsis definitions with microbiological characteristics in patients admitted through a sepsis alert system; a prospective cohort study. Annals Clin. Microbiol. Antimicrob. 21, 7 (2022).

14. Huerta, L. E. & Rice, T. W. Pathologic difference between sepsis and bloodstream infections. The journal applied laboratory medicine 3, 654–663 (2019).

15. Bates, D. W. et al. Predicting bacteremia in patients with sepsis syndrome. J. Infect. Dis. 176, 1538–1551 (1997).

16. Tjandra, K. C. et al. Diagnosis of bloodstream infections: an evolution of technologies towards accurate and rapid identification and antibiotic susceptibility testing. Antibiotics 11, 511 (2022).

17. PTopić opović, N., Kazazić, S. P., Bojanić, K., Strunjak-Perović, I. & Čož-Rakovac, R. Sample preparation and culture condition effects on maldi-tof ms identification of bacteria: A review. Mass spectrometry reviews 42, 1589–1603 (2023).

18. Perše, G. et al. Sepsityper® kit versus in-house method in rapid identification of bacteria from positive blood cultures by maldi-tof mass spectrometry. Life 12, 1744 (2022).

19. Chen, J. H. et al. Direct bacterial identification in positive blood cultures by use of two commercial matrix-assisted laser desorption ionization–time of flight mass spectrometry systems. J. clinical microbiology 51, 1733–1739 (2013).

20. Buchan, B. et al. Clinical evaluation of the accelerate arc module and bc kit for isolation of microorganisms from positive blood culture broths and suitability for maldi-tof analysis. In JOURNAL OF MOLECULAR DIAGNOSTICS vol. 24 S51–S51 (ELSEVIER SCIENCE INC STE 800, 230 PARK AVE, NEW YORK, NY 10169 USA, 2022).

21. Gajic, I. et al. Antimicrobial susceptibility testing: a comprehensive review of currently used methods. Antibiotics 11, 427 (2022).

22. Baltekin, Ö., Boucharin, A., Tano, E., Andersson, D. I. & Elf, J. Antibiotic susceptibility testing in less than 30 min using direct single-cell imaging. Proc. Natl. Acad. Sci. 114, 9170–9175 (2017).

23. Osaid, M. et al. A multiplexed nanoliter array-based microfluidic platform for quick, automatic antimicrobial susceptibility testing. Lab on a Chip 21, 2223–2231 (2021).

24. Pitt, W. G. et al. Rapid separation of bacteria from blood—review and outlook. Biotechnol. progress 32, 823–839 (2016).

25. Miguelez, M. H. M. et al. Culture-free rapid isolation and detection of bacteria from whole blood at clinically relevant concentrations. bioRxiv 2024–05 (2024).

26. Forsyth, B. et al. A rapid single-cell antimicrobial susceptibility testing workflow for bloodstream infections. Biosensors 11, 288 (2021).

27. Zeng, K., Osaid, M. & van der Wijngaart, W. Efficient filter-in-centrifuge separation of low-concentration bacteria from blood. Lab on a Chip 23, 4334–4342 (2023).

28. Kim, T. H. et al. Blood culture-free ultra-rapid antimicrobial susceptibility testing. Nature 1–10 (2024).

29. Ohlsson, P. et al. Integrated acoustic separation, enrichment, and microchip polymerase chain reaction detection of bacteria from blood for rapid sepsis diagnostics. Anal. chemistry 88, 9403–9411 (2016).

30. Mach, A. J. & Di Carlo, D. Continuous scalable blood filtration device using inertial microfluidics. Biotechnol. bioengineering 107, 302–311 (2010).

31. Narayana Iyengar, S., Kumar, T., Mårtensson, G. & Russom, A. High resolution and rapid separation of bacteria from blood using elasto-inertial microfluidics. Electrophoresis 42, 2538–2551 (2021).

32. Li, S. et al. Acoustofluidic bacteria separation. J. Micromechanics Microengineering 27, 015031 (2016).

33. Park, S., Zhang, Y., Wang, T.-H. & Yang, S. Continuous dielectrophoretic bacterial separation and concentration from physiological media of high conductivity. Lab on a Chip 11, 2893–2900 (2011).

34. Kayin, M., Mert, B., Aydemir, S. & Özenci, V. Comparison of rapid bacpro® ii, sepsityper® kit and in-house preparation methods for direct identification of bacteria from blood cultures by maldi-tof ms with and without sepsityper® module analysis. Eur. J. Clin. Microbiol. & Infect. Dis. 38, 2133–2143 (2019).

35. Croxatto, A., Prod’hom, G., Durussel, C. & Greub, G. Preparation of a blood culture pellet for rapid bacterial identification and antibiotic susceptibility testing. J. visualized experiments: JoVE 51985 (2014).

36. Kinahan, D. J. et al. Density-gradient mediated band extraction of leukocytes from whole blood using centrifugo-pneumatic siphon valving on centrifugal microfluidic discs. PloS one 11, e0155545 (2016).

37. Kandavalli, V., Karempudi, P., Larsson, J. & Elf, J. Rapid antibiotic susceptibility testing and species identification for mixed samples. Nat. Commun. 13, 6215 (2022).

38. Camsund, D. et al. Time-resolved imaging-based crispri screening. Nat. methods 17, 86–92 (2020).

